# Long-Term Neurovascular Dysfunction and Altered Placenta-Derived EV miRNA Profiles in a Rat Model with Preeclamptic Features

**DOI:** 10.1101/2025.09.22.675363

**Authors:** Jonathan Ray, Brian M. Westwood, Xuming Sun, Robert Gould, Yixin Su, Marli Graves, Nildris Cruz-Diaz, Sangeeta Singh, Ashish Kumar, Gagan Deep, Hossam A. Shaltout, NyKea L. Lewis, Tao Ma, Jeffrey Denney, Victor M. Pulgar, Liliya M. Yamaleyeva

## Abstract

**Objective:** Preeclampsia increases long-term cardiovascular, cerebrovascular, and cognitive risk but the underlying mechanisms remain unclear. We examined cerebrovascular function, cognitive performance, and placenta-derived extracellular-vesicle (PD-EV) microRNA profiles in a transgenic rat model with preeclamptic features (TGAPE) at 2 months after delivery (2M).

**Methods:** Middle cerebral artery (MCA) hemodynamics were assessed by transcranial Doppler (TCD), and cerebral oxygenation was measured by photoacoustic imaging (PAI) in TGA-PE and Sprague-Dawley control rats. Blood pressure was measured using a tail-cuff method. Touchscreen-based cognitive tasks evaluated learning and reward-directed responding, while histological assessment evaluated MCA structure. MicroRNA sequencing was performed on circulating PD-EV, and miRNA target networks were analyzed using KEGG-based NEST-Scoring (Node, Edge, System, Topology). Validation of miR-363-3p was performed in PD-EV, plasma, non-PD-EV, and human brain endothelial cells (HBEC-5i).

**Results:** TGA-PE rats were hypertensive and exhibited lower MCA resistance and pulsatility indexes with unchanged blood flow velocities. Cognitive testing revealed impaired associative learning and reduced motivation to respond on a progressive-ratio schedule. Temporal brain oxygen saturation was higher in TGA-PE rats with unchanged total hemoglobin, lower brain HIF-2α, and unchanged VEGF levels. MCA elastin content was reduced in TGA-PE rats without changes in total collagen. PD-EV sequencing identified 29 differentially expressed miRNAs, including reduced miR-363-3p in non-placental EVs and plasma at 2M. Inhibition of miR-363-3p in human cerebral endothelial cell line HBEC-5i upregulated NOX4, p21, and β-galactosidase levels. NEST-Scoring highlighted pathways involving MAPK, Ras, TNF, focal adhesion, cytoskeletal regulation, and cellular senescence.

**Conclusions:** The exposure to experimental preeclampsia resulted in altered MCA flow resistance, cerebral oxygenation, vascular structure and behavioral performance. Although direct mechanistic studies are needed to establish the role of miRNA363-3p in neurovascular health, the increased endothelial oxidative stress and senescence-associated markers in HBEC-5i cells in response to miRNA363-3p inhibition suggest a potential protective role for miRNA363-3p. A novel NEST-Scoring analysis also identified potential pathways for future mechanistic investigations. Overall, the TGA-PE model provides a useful experimental system for investigating candidate mechanisms that may contribute to long-term maternal neurovascular changes following hypertensive pregnancy and sequelae of preeclamptic pathophysiology.

**Graphical Abstract:** Long-Term Neurovascular Dysfunction and Altered Placental EV miRNA Profiles in a Rat Model with Preeclamptic Features. TGA-PE rats exhibited long-term cerebrovascular remodeling characterized by altered cerebral hemodynamics and vascular structure, accompanied by long-term cognitive dysfunction. A systemic reduction of miR-363-3p was observed in extracellular vesicles and plasma. In human cerebral endothelial cells, miR-363-3p inhibition promoted oxidative stress and endothelial senescence. These findings support the TGA-PE model as a translational platform for investigating mechanisms underlying persistent cerebrovascular dysfunction associated with preeclampsia. (generated with Copilot Opus 5.5)

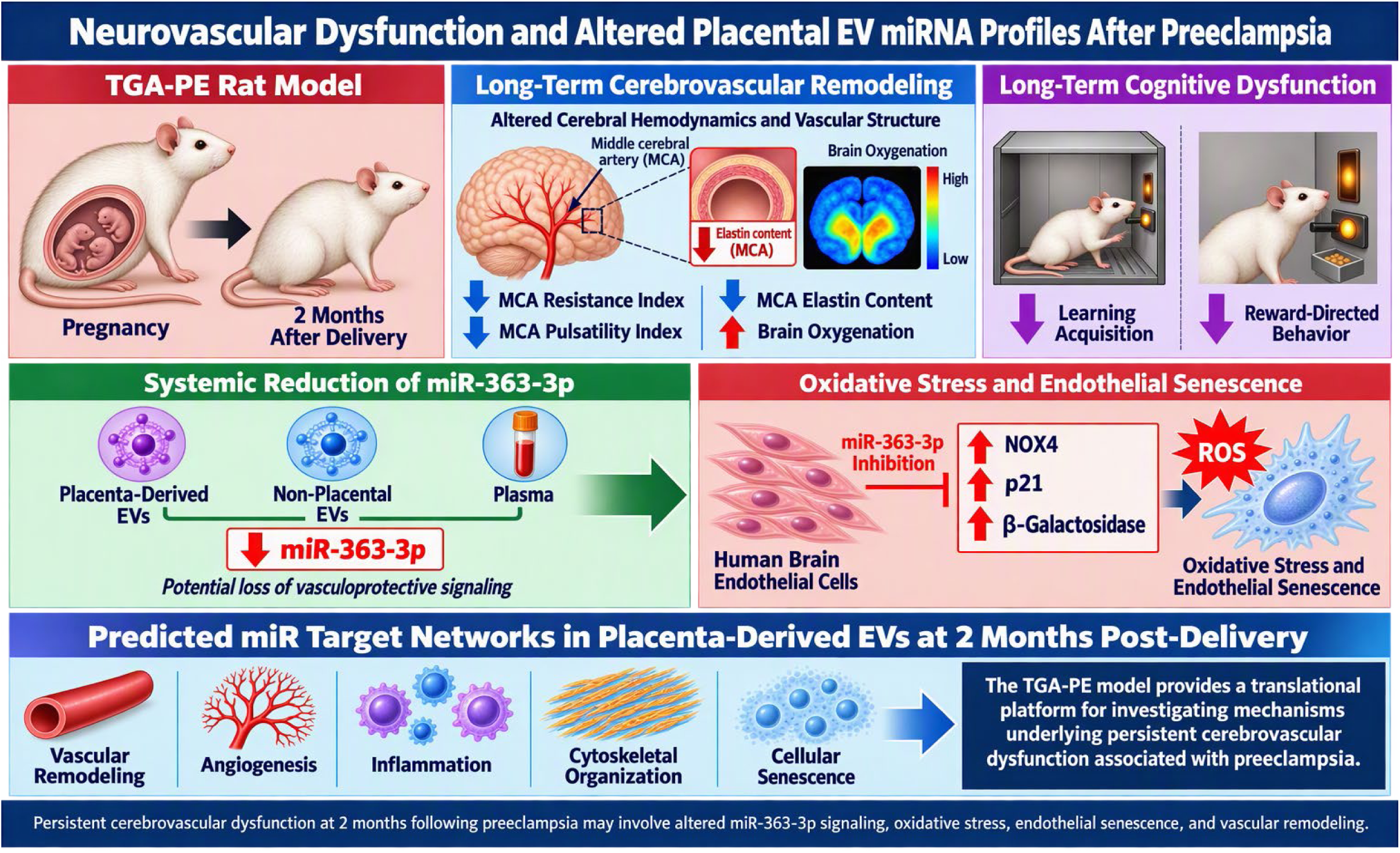

## INTRODUCTION

Preeclampsia is a multisystem hypertensive disorder of pregnancy with heterogeneous clinical presentation and a major contributor to maternal morbidity and mortality [1, 2]. Beyond its acute manifestations, it is associated with increased risk of hypertension, cerebrovascular events, stroke, vascular dementia, and early cognitive impairment [3–8]. Women with a history of preeclampsia also exhibit a higher risk of developing symptomatic cognitive impairment compared to those without preeclampsia [3]. Importantly, cerebrovascular and neurological complications may manifest during pregnancy or emerge years to decades after delivery. Accordingly, preeclampsia is now recognized by the American Heart Association and American Stroke Association as an independent risk factor for stroke in women, underscoring the need for long-term cerebrovascular surveillance in this population [9, 10].

Transcranial Doppler ultrasound (TCD) is a validated, non-invasive diagnostic tool used to assess hemodynamics of cranial vessels, including the middle cerebral artery (MCA), which supplies approximately two-thirds of the cerebral blood flow. Clinical studies demonstrate that postmenopausal women with prior preeclampsia exhibit lower resting MCA blood flow velocity and reduced cerebrovascular reactivity to hypercapnia compared with women with normotensive pregnancies, indicating persistent impairment of vascular responsiveness in later life [7, 11]. These cerebrovascular changes may have clinical relevance as women with a history of preeclampsia show poorer cognitive performance, particularly in executive function and information-processing speed domains typically associated with vascular cognitive impairment with deficits observed from midlife through older age, or at least 19 years after delivery [4, 12, 13]. Together, these findings implicate cerebrovascular disease as an important pathway to cognitive impairment following preeclampsia; however, the mechanisms linking preeclampsia to persistent maternal neurovascular abnormalities remain unclear.

Experimental models support a vascular contribution to the long-term outcomes in preeclampsia, though most studies have focused on changes during gestation or the immediate postpartum period, with fewer addressing the later post-preeclampsia period. Placental ischemia during pregnancy impairs MCA myogenic responsiveness, causes loss of cerebral blood flow autoregulation, increased blood-brain barrier permeability, and cerebral edema during pregnancy - providing mechanistic evidence that preeclampsia disrupts cerebrovascular hemodynamics [14, 15]. Notably, persistent postpartum neuroinflammation, reduced occludin expression, activated microglia, and regional cerebral edema were reported in the reduced uterine perfusion pressure (RUPP) model of preeclampsia at two months after delivery, indicating that placental ischemia is a driver of prolonged cerebrovascular injury [16]. Nevertheless, the circulating molecular signals associated with later cerebrovascular and cognitive abnormalities have not been fully characterized.

Although the pathogenesis of preeclampsia is multifactorial, placental malperfusion promotes the release of soluble factors into the maternal circulation which are thought to play important roles in mediating maternal endorgan damage, including cerebrovascular dysfunction [7, 17–19]. The placenta communicates with maternal systems through soluble factors, cytokines, and extracellular vesicles (EVs). Placenta-derived EVs (PD-EVs) and their cargo particularly microRNAs (miRNAs) have emerged as candidate mediators capable of directly or indirectly influencing maternal cerebrovascular function through interactions with the blood-brain barrier or the neurovascular unit [20, 21]. These EV-associated molecular signals may provide mechanistic insight into how placental ischemia/hypoxia in preeclampsia drive long-term cerebrovascular complications. A better understanding of placenta-derived molecular signals could improve postpartum risk stratification and inform preventive surveillance strategies for women at risk for long-term complications following preeclampsia [22]. Moreover, placental dysfunction has been linked to alterations in junction protein regulation and dopamine receptor function that may affect maternal vascular and behavioral outcomes [23, 24]. In addition to these placenta-centered mechanisms, women with a history of preeclampsia exhibit accelerated biological aging profiles that may amplify long-term cardiovascular and cerebrovascular risk [25, 26].

We used a transgenic rat model generated by mating female rats expressing the human angiotensinogen gene with males expressing the human renin gene [27–29]. This TGA-PE model reproduces major features of human preeclampsia including hypertension, proteinuria, and cardiovascular dysfunction [30–33]. Given the critical role of PD-EVs in intercellular signaling, we hypothesized that placentally-derived EV signatures including miRNAs and their associated target gene pathways would be detectable and differentially expressed at term and postpartum. To test this hypothesis, we performed miRNA sequencing of circulating PD-EVs isolated from preeclamptic transgenic rats and normotensive controls at late gestation and two months post-delivery - a time corresponding to approximately 4-5 human years post-pregnancy [34]. The differences in miRNA profiles may help identifying molecular targets associated with long-term cerebrovascular dysfunction and cognitive impairment for future postpartum therapeutic intervention.

Furthermore, to better characterize pathway-level alterations in molecular signaling, we developed and applied a novel network analytical method - NEST-Scoring (Node, Edge, System, Topology) - a pathway-based approach that evaluates the distribution and clustering of miRNA target genes within biological networks [35–39]. To grasp the nuances of a network and the related distances between elements, NEST-Scoring provides a graphical framework for assessing whether signaling patterns are diffused or concentrated across a pathway. By emphasizing first-neighbor relationships among genes of interest, this approach is particularly suited for exploratory pathway organization analyses in the context of EV-associated miRNA target networks. It weighs each node by its connectivity to other genes of interest (excitatory influence) relative to its connectivity to genes outside that set (inhibitory influence) [40]. We therefore applied NEST-Scoring to evaluate whether PD-EV miRNA target gene networks differ between normotensive and preeclamptic rats at both late gestation and the two-month post-delivery time point.

The molecular analysis of circulating PD-EVs was complemented by a multi-scale physiological approach integrating: (i) TCD assessment of MCA hemodynamics, (ii) photoacoustic imaging of MCA and adjacent brain oxygenation, and (iii) cognitive function testing using touchscreen-fitted operant chambers. Accordingly, the specific objectives of this study were to: 1) determine whether MCA hemodynamics, brain oxygenation, and cognitive function are altered in transgenic rats with preeclamptic features at two months post-delivery; and 2) identify whether miRNA profiles of circulating PD-EVs and associated target gene pathways assessed using KEGG-based NEST-Scoring network analysis, reveal novel regulatory pathways relevant to the long-term cerebrovascular dysfunction observed after preeclampsia.

## METHODS

### Animals

The Institutional Animal Care and Use Committee of Wake Forest University School of Medicine approved the study (A21-057), and all experiments were conducted in accordance with relevant guidelines and regulations. Female transgenic rats overexpressing human angiotensinogen (TGA) and male transgenic rats overexpressing human renin (TGR) were obtained from the Department of Surgery/Hypertension and Vascular Research Center Transgenic Core at Wake Forest University School of Medicine, Winston-Salem, North Carolina. Sprague-Dawley (SD) rats purchased from Charles River Laboratories (Wilmington, MA, USA) were used as controls. All rats were housed under controlled temperature, humidity, and a 12:12-h light-dark cycle, fed standard rodent chow (Lab Diet 5P00–Prolab RMH 3000, PMI Nutrition International, Inc., Brentwood, MO), and provided water *ad libitum* throughout the experimental protocols.

### Study timeline

Female TGA rats were mated with male TGR rats to induce preeclampsia-like phenotype (TGA-PE), and SD females were mated with SD males. TGA-PE and SD rats aged 11–26 weeks during late gestation were used in this study. Gestational day (GD) was estimated by embryonic features detected by transabdominal ultrasound [41]. Plasma was collected from SD and TGA-PE rats during late gestation (LG) and at 2 months post-pregnancy (2M). Blood pressure, heart rate, and transcranial Doppler (TCD) ultrasound analysis of the left middle cerebral artery (LMCA) were performed during late gestation (GD17–19) and at 2M.

### Blood pressure and heart rate measurements

Systolic, diastolic, and mean blood pressures and heart rate were measured in trained, conscious rats by tail-cuff plethysmography using the Non-Invasive Blood Pressure Monitoring System (Columbus Instruments, Columbus, OH) one day before TCD ultrasound as previously described [42]. At least ten blood pressure measurements were recorded, and values were averaged for each animal. The data were reported as mean ± SEM.

### Transcranial Doppler ultrasound

High-frequency ultrasound was used to measure pulsatility index (PI) and resistance index (RI) of the LMCA. Rats were placed on a temperature-controlled platform, and temporal hair was removed using depilatory cream (Nair, Church & Dwight Co., Ewing, NJ). Ultrasound imaging was performed under 1.5% isoflurane anesthesia using a Vevo LAZR Ultrasound and Photoacoustic System with an LZ250 transducer (FujiFilm VisualSonics, Toronto, Canada). The LMCA was visualized in color Doppler mode by directing the transducer through the temporal approach as previously described [43]. Peak systolic (V_max_), minimum diastolic (V_min_), and mean (V_mean_) blood flow velocities were measured by pulse-wave Doppler and averaged over three cardiac cycles. PI and RI were calculated as follows: PI = (V_max_ − V_min_)/V_mean_; RI = (V_max_ − V_min_)/V_max_. Data were analyzed using Vevo LAB software version 5.7.1 for Windows (FujiFilm VisualSonics, Toronto, Canada).

### Ultrasound-guided photoacoustic imaging for in vivo measurements of MCA and brain oxygenation (temporal lobe)

Rats were anesthetized with 1.5% isoflurane and placed on a heated platform for photoacoustic imaging (PAI). Heart rate, measured by electrocardiogram electrodes connected to the platform, was on average at 350– 400 beats/min. Temporal hair was removed using depilatory cream (Nair, Church & Dwight Co., Inc.). Imaging was performed using a Vevo LAZR system with an LZ250 transducer operating at 13–24 MHz and providing an axial resolution of approximately 75 µm. The integrated optical component provided tunable laser excitation (680–970 nm) with peak optical fluence up to 20 mJ/cm² in compliance with American National Standards Institute (ANSI) safety guidelines [43]. Vevo LAZR software tools (HemoMeaZure and OxyZated) co-registered the photoacoustic signal from hemoglobin with real-time anatomic tissue images and calculated mean sO_2_ concentration. Imaging parameters were standardized across experiments to minimize variability, including background noise reduction, consistent photoacoustic signal amplitude settings, and a fixed imaging depth for all animals.

### Cognitive function

SD and TGA-PE rats were trained to respond on a touch-sensitive computer screen (Lafayette Instruments, Lafayette, IN) whereby correct responses resulted in delivery of a single 45-mg sucrose pellet. To assess associative learning, rats completed a pairwise discrimination task by learning to discriminate between two stimuli presented pseudo-randomly on the screen in daily cognition sessions. After completion, rats were trained to respond on a single stimulus for sucrose pellets; the number of responses on the stimulus increased following each subsequent completed ratio. This Progressive Ratio task (PR) served as a measure of motivated responding.

### Total collagen and elastin content detection in MCA

Total collagen and elastic fibers in the MCA of TGA-PE and SD rats were assessed using Trichrome or Verhoeff’s staining, respectively (American MasterTech; Cat#: KTVEL). The Trichrome-or Verhoeff-positive area was normalized to the total vessel area. Signal intensity in the MCA media was normalized to background intensity and to the maximum RGB value of 255, and results were reported as relative intensity units using Image J software as described by us [44].

### Immunohistochemistry of MCA and brain for HIF2α and VEGF

LMCAs and brains were fixed in 10% formalin, embedded in paraffin, and cut into 5-μm sections. Immunostaining was performed using the avidin biotin complex (ABC) method with a diaminobenzene solution used as the chromogen. Antigen retrieval treatment with IHC-TEK Epitope Retrieval Solution (IHC World, Woodstock, MD, USA) was applied at 95-98°C for 40 min. Non-specific binding was blocked in a buffer containing 10% normal goat serum, 0.1% bovine serum albumin, and 1% Triton X-100 in PBS for 30 min. LMCA sections were incubated with rabbit polyclonal HIF2a antibody (1:300 dilution; Abcam Inc., Waltham, MA, USA; cat No. ab199), rabbit polyclonal VEGF antibody (1:300 dilution; Santa Cruz Biotechnology, Inc., Dallas, TX, USA; cat No. sc-507), and secondary biotinylated goat anti-rabbit antibodies (1:500 dilution; Vector Laboratories Inc., Newark, CA, USA; cat No. BA-1000). The images were acquired with a NanoZoomer^®^ S360MD Slide scanner system (Hamamatsu Photonics K.K., Hamamatsu City, Shizuoka Prefecture, Japan) at up to 20x mode and analyzed by Image J software as described by us [44].

### Cell culture

HBEC-5i, a human cerebral microvascular endothelial cell line was purchased from The American Type Culture Collection (ATCC, Manassas, Virginia, USA; cat No. CRL-3245). The cell line was maintained in DMEM/F12 (Thermo Fisher Scientific Inc., Waltham, MA, USA; cat. No. 11330057) containing 10% fetal bovine serum (FBS; Thermo Fisher Scientific Inc., Waltham, MA, USA; cat No. 26140079) and 40mg/ml Endothelial Cell Growth Supplement (ECGS; Sigma-Aldrich, St. Louis, MO, USA; cat. No. E0760-15mg). HBEC-5i cells at a passage 3-4 were seeded at a density of 3 x10^5^ cells per 60 mm dish coated with 0.1% Gelatin (Sigma-Aldrich, St. Louis, MO, USA; cat. No. ES-006-B). When cells reached 60-80% confluency, miRCURY LNA^TM^ has-mir-363-3P power inhibitor (QIAGEN, Hilden, Germany; cat No. 339131 YI04101050-DDB) or miRCURY LNA^TM^ miRNA power inhibitor negative control A (QIAGEN, Hilden, Germany; cat No. 339136 YI00199006-DDB) was added at 50 nM concentration for 48 hours. After collection of the conditioned cell culture media, the cells were washed with 1X Gibco dPBS (Fisher Scientific, Waltham, MA, USA; cat No. 14190144) and collected after 20-minute incubation on ice with lysis buffer containing 10 mM Hepes/125 mM NaCl, pH 7.4, 10 µL/mL protease inhibitor cocktail (Sigma-Aldrich, St. Louis, MO, USA; cat. No. P8340), 10 µL/mL phosphatase inhibitor cocktail 2 (Sigma-Aldrich, St. Louis, MO, USA; cat. No. P5726), 1 mM EDTA, and 1% Triton X-100.

### Western blotting

The cell lysate samples were subjected to 3-second sonication pulses at 20 kHz for 15 seconds. Samples were then centrifuged (10,000 g, 5 min, 4℃) to remove debris. Total protein levels in the supernatant were determined using the Pierce^TM^ BCA Protein assay kit (Thermo Fisher Scientific Inc., Waltham, MA, USA; cat No. 23227) according to the manufacturer’s protocol. 5µg of protein from each sample were separated by 4-20% TGX Stain-Free gel (Bio-Rad Laboratories, Hercules, CA, USA; cat No. 567-8095) and transferred onto a PVDF membrane with the Trans-Blot Turbo Transfer System (Bio-Rad Laboratories, Hercules, CA, USA). The membrane was blocked with 5% non-fat milk or BSA in Tris-buffered saline with 0.05% Tween (blocking buffer) for 1 hour at room temperature, then incubated with primary antibodies in the blocking buffer overnight at 4°C. Following incubation with the corresponding horseradish peroxidase-conjugated secondary antibody at room temperature for 1 hour, protein bands were visualized by using SuperSignal^TM^ West Femto Kit (Thermo Fisher Scientific Inc., Waltham, MA, USA; cat No. 34095). The images were captured using a ChemiDoc Imaging System (Bio-Rad Laboratories, Hercules, CA, USA) and analyzed using Image Lab software (Bio-Rad Laboratories, Hercules, CA, USA). Each band was expressed in arbitrary units and normalized to β-actin. The primary antibodies used were rabbit anti-β-actin antibody (1:1000 dilution; Cell Signaling Technology, Danvers, MA, USA; cat No. 8457); rabbit anti-Beta Galactosidase antibody (1:500 dilution; Proteintech Group Inc., Rosemont IL, USA; cat No. 15518-1-AP); rabbit anti-p21 antibody (1:500 dilution; Thermo Fisher Scientific Inc., Waltham, MA, USA; cat No. MA5-42680); rabbit anti-NOX4 antibody (1:500 dilution; Abcam Inc., Waltham, MA, USA; cat No. ab133303).

### Isolation of PD-EV

Total small EV were isolated from plasma (n=3 from each group) via sequential centrifugation and ExoQuick precipitation as reported by us previously [45–47]. Total EVs were characterized by nanoparticle tracking analyses (NTA) using Nanosight NS300 (Malvern Instruments, UK) for size and concentration as reported by us earlier [47, 48]. Next, 2,500 μg of total EVs were incubated with 5 μg of biotin-tagged PLAP antibody (MA5-38111; ThermoFisher) overnight at 4°C with constant mixing. PLAP antibody was labeled with biotin using FluoReporter Mini-Biotin-XX protein labeling Kit (ThermoFisher). Streptavidin-tagged magnetic beads were added to the samples and incubated for 2 hrs at RT with continuous mixing. Beads were then washed twice with 0.1% TBST and PLAP+ PD-EV were eluted from the beads by incubating with 200 μl elution buffer for 10 min at RT. Beads were then magnetized and supernatant with PD-EV was collected in 10% v/v 1M Tris buffer (pH 9) to neutralize the pH.

### Nano-flow cytometry

Total EVs were analyzed using nano-flow cytometry following the method described by us previously [46]. Briefly, total EVs were incubated with fluorescently labeled PLAP antibody (ThermoFisher, Cat. No. MA5-38111), for 2 h at room temperature in the dark. PLAP antibody was fluorescently labeled using APC conjugation kit (Abcam, Cat. No. ab201807) as per the manufacturer’s recommendations. Following antibody incubation, membrane-labeling dye CellBrite steady 488 (CellBrite steady membrane staining Kit, Biotium, Fremont, CA, USA) was diluted 200-fold in 0.1 micron filtered PBS, and 50 uL of diluted dye was added to the total EVs for 15 min incubation in the dark. Samples were diluted 100-to 200-fold in 0.1% BSA in PBS filtered through a 0.1-micron filter to achieve an aborted ratio of less than 10%. Total EVs were acquired on CytoFlex (Beckman Coulter Life Science, Indianapolis, United States) for 60s at a low flow rate. Filtered PBS was run for 60s in between the samples. Total EVs without dye was used to set the gate for dye-positive events, and total EVs labeled with dye but without fluorescent antibody were used to set the gate for AF647-positive events.

### PD-EV RNA for sequencing analysis

Total RNA from PD-EV was isolated using Trizol based method as detailed previously [47]. Briefly, 200 μl of PD-EV were incubated with 20 μl RNAse A for 30 min at RT to remove any non-EV RNA. Next, 800 μl TRIZol was added to the samples and incubated at RT for 10 min with occasional vortexing in between. Following incubation, 150 μl of chloroform was added and samples were incubated for 2 min before centrifugation at 14,000g for 15 min at 4°C. The top clear phase was collected and mixed with 2 volumes of ethanol for overnight incubation at -20°C. Afterwards samples were purified using RNA isolation columns (Qiagen, Germantown, Maryland, USA), following kit’s instruction, and finally eluted in RNAse free water. The concentration of RNA was measured by NanoDrop (ThermoFisher Scientific). Small RNA sequencing using 350 ng of PD-EV RNA was performed by LC Sciences (Houston, TX, USA).

### Plasma miRNA isolation and quantitative real-time RT-PCR

All kits, primers, and reagents were purchased from QIAGEN (Hilden, Germany). Total RNA, including miRNA, was isolated from 0.2 ml of rat plasma using the QIAGEN miRNeasy Serum/Plasma Kit (cat no. 217184) according to the manufacturer’s protocol. Total RNA concentration and purity were measured using a NanoDrop^TM^ One^C^ Microvolume UV-Vis Spectrophotometer (ThermoFisher Scientific Inc., Waltham, MA). First-strand cDNA was synthesized from 1 µg total RNA using the miRCURY LNA RT Kit (cat no. 339340). Quantitative real-time RT-PCR was performed on a QuantStudio 3 Real-Time PCR System (Applied Biosystems, Foster City, CA) using the miRCURY miRNA SYBR Green PCR Kit (cat no. 339345, QIAGEN). Gene-specific primers included rno-miR-363-3p (339306-YP02103062) and 5S rRNA (339306-YP00203906). Each 10-µl reaction contained 3 µl of 1:10 diluted first-strand cDNA, 5 µl of 2x miRCURY SYBR Green Mastermix, 1 µl of forward and reverse primer mix, 0.05 µl ROX reference dye, and 0.95 µl RNase-free water. The PCR protocol consisted of 95°C for 2 min, followed by 40 cycles of 95°C for 10 s and 60°C for 60 s, with a melting curve stage of 95°C for 15 s, 60°C for 60 s, and 95°C for 1 s. All samples were run in duplicate, and 5S rRNA was used as the reference gene. Relative target miRNA levels were normalized to 5S rRNA in the same sample using the 2^-ΔΔCt^ method (Ct = threshold cycle) and reported relative to the mean control value.

### Statistics

Group comparisons were performed using two-way analysis of variance (ANOVA) followed by Bonferroni’s post hoc test or by unpaired t-test when appropriate (GraphPad Software, San Diego, CA). Values of p < 0.05 were considered statistically significant. All data are presented as mean ± SEM.

### Discrete correlate summation

Top-ranked miRNAs identified by IQR-score [35] were clustered using discrete correlate summation (DCS) [49]. Target gene sets were then assessed for functional enrichment using DAVID [50, 51] and KEGG pathway analyses [52] (Supplemental Figures 1-2). miRNA targets were stratified by miRDB target score (MTS) [53]. High-confidence targets (MTS ≥ 95) related to blood pressure and pregnancy, based on Swiss-Prot annotations [54], were selected (n = 146; “95BpPreg”) and compared with general high-confidence targets (MTS ≥ 80; n = 672; “MTS80”; Figure 1).

**Figure 1.**
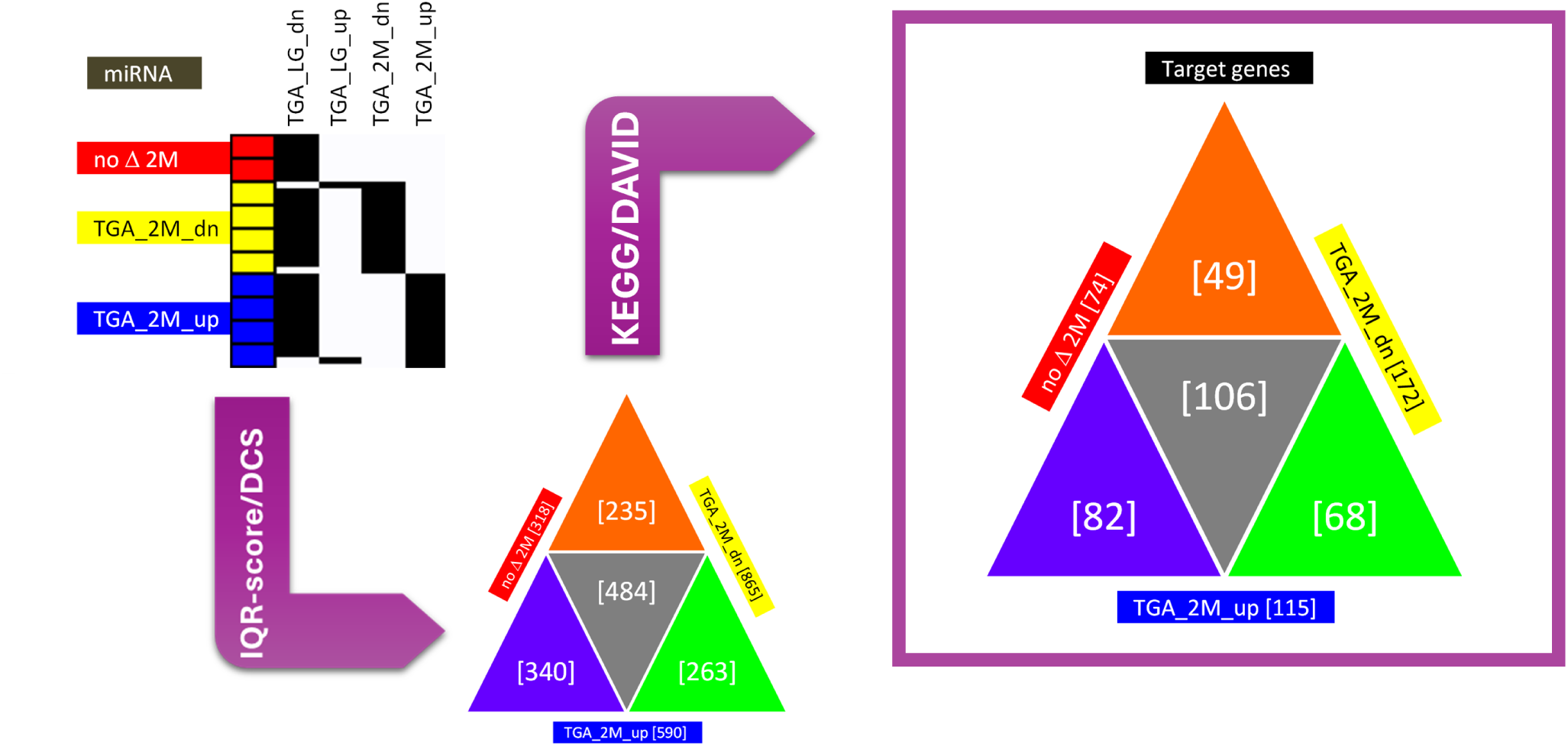
PD-EV miRNA target gene enrichment pipeline for NEST-Scoring evaluation. Triangular compositional diagram output (on right, outlined) of the gene targets for the three broad groups of miRNAs [no Δ 2M, TGA 2M down, TGA 2M up] after IQR-score/DCS clustering and KEGG pathway gene enrichment analysis in DAVID (see Supplemental Table Figure 1).

### NEST-Scoring

53 KEGG pathways NEST-Scored (Figure 2) for miRDB targets score (MTS)≥95 (yellow nodes) and Swiss-Prot blood pressure (cyan nodes) and pregnancy (magenta nodes) MTS≥80 genes relative to all other IQR-score/DCS MTS80 gene targets (gray nodes) (Figure 2). The NEST-Score is simply a scoring metric to rank KEGG pathway networks, utilizing the total MTS ≥ 95 (+1; blue cross, counter 5) and BpPreg (+1; black cross, counter 7) node counts relative to whether the nodes are ‘nested’ near MTS ≥ 80 nodes (+0.25; cyan cross, counter 2)/each edge vs. near other rat genome nodes (-0.25; gray cross, counter 9)/each edge.

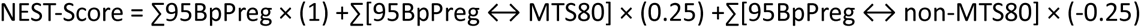

**Figure 2.**
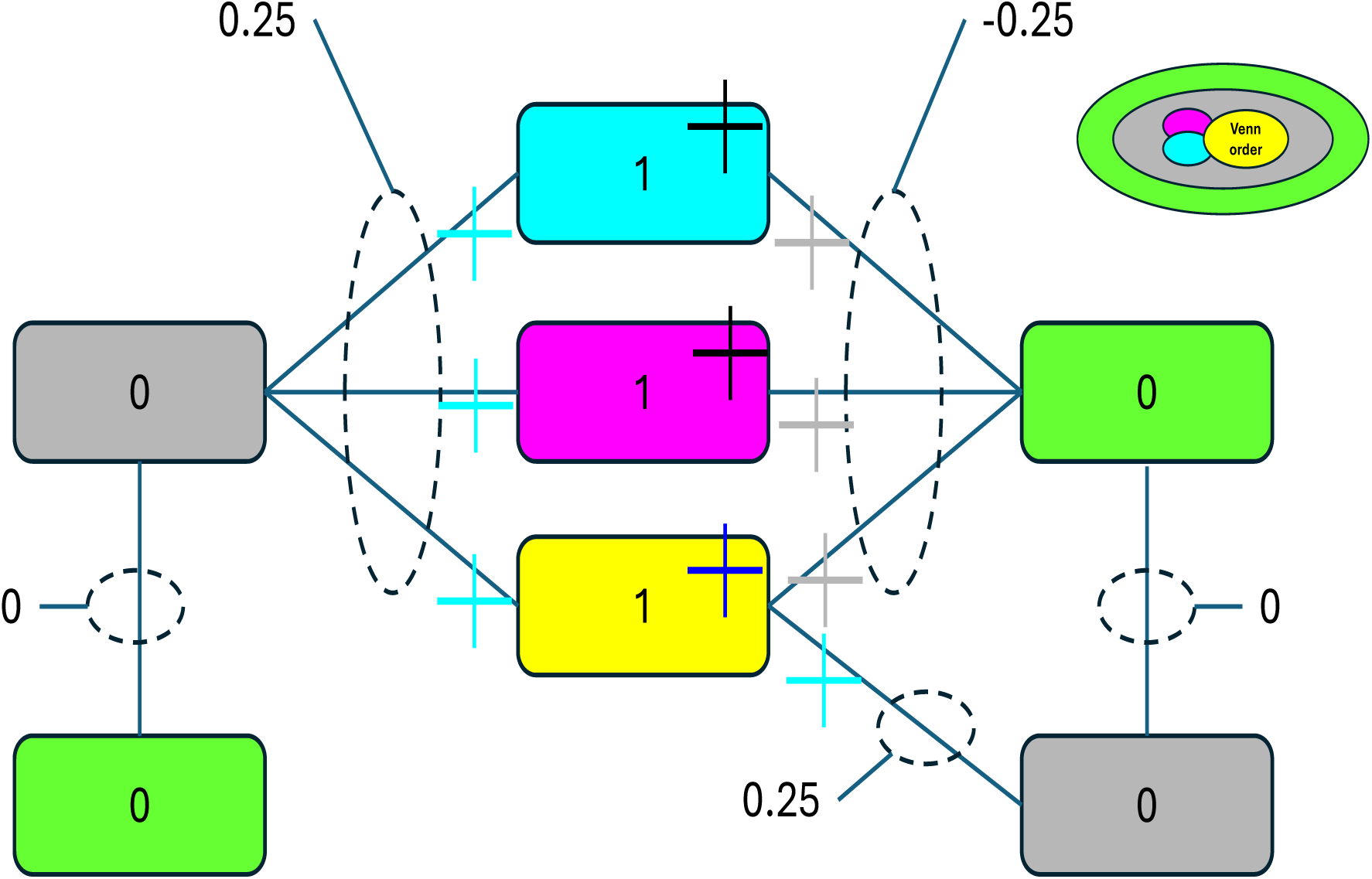
Cartoon example of NEST-Scoring evaluation of miRNA used for 53 KEGG pathways. Nodes in KEGG pathways are yellow-95 if miRDB score >95, masked cyan-Bp, magenta-Preg, otherwise gray (green are rat genome). Scor-ing: CMY nodes=1, edge from CMY-gray = 0.25 / CMY-green = -0.25.

## RESULTS

Systolic and diastolic blood pressures were higher in TGA-PE rats than in SD rats during late gestation and at 2M (Figure 3A,B). In TGA-PE rats, blood pressure decreased at 2M compared with late gestation, whereas systolic and diastolic blood pressures did not differ over time in SD rats. Heart rate was higher in TGA-PE rats during late gestation but did not differ between TGA-PE and SD rats at 2M (Figure 3C). Heart rate also decreased in TGA-PE rats at 2M compared with late gestation (Figure 3C). We have previously reported that TGA-PE rats have lower body weight vs. SD controls [32]. No differences in body weight were recorded between TGA-PE and SD females at 2M (Figure 3D).

**Figure 3.**
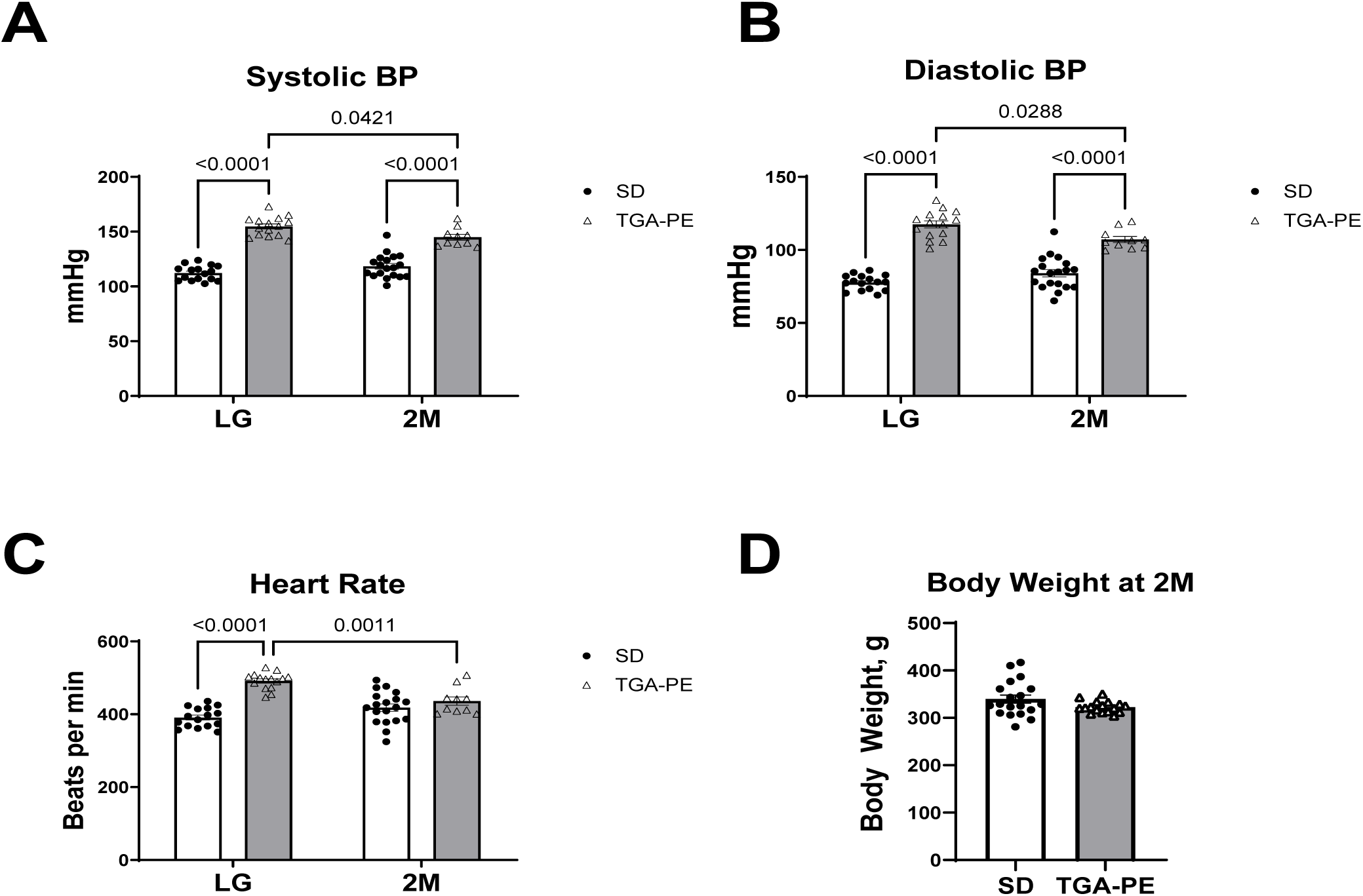
Blood pressure, heart rate and body weight in TGA-PE vs. SD rats at LG and 2M post-delivery (2M). Data are mean ± SEM, n=10-19.

Peak systolic, minimum diastolic, and mean LMCA velocities did not differ between strains or time points although there was a trend for an increase of LMCA minimum diastolic velocity in TGA-PE at LG (p=0.05) (Figure 4A,B). However, LMCA resistance and pulsatility indexes were lower in TGA-PE rats than in SD rats at 2 months post-delivery (Figure 4C).

**Figure 4.**
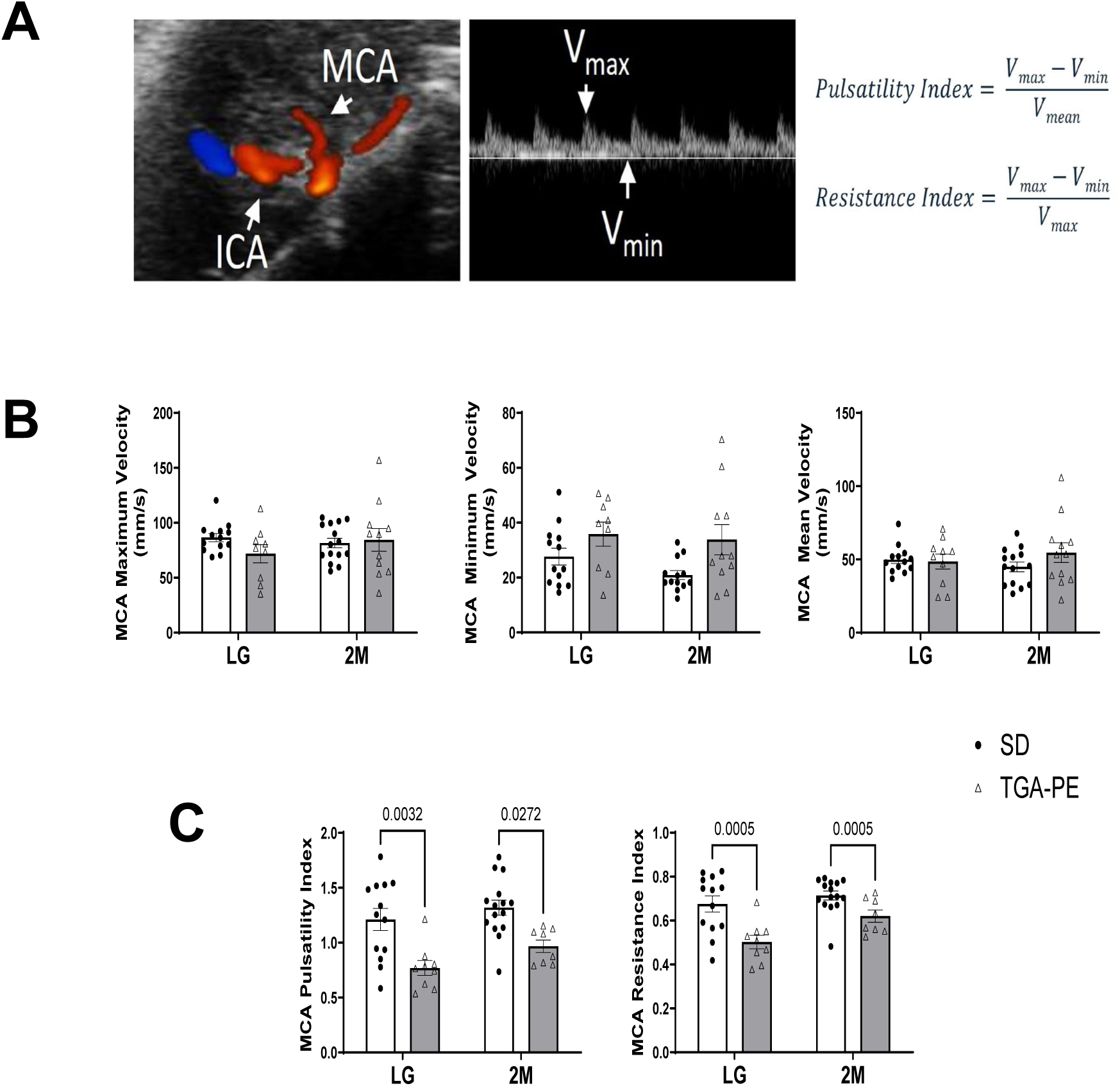
MCA velocity and resistance in TGA-PE vs. SD rats at late gestation (LG) and two months (2M) post-delivery. Panel A shows a representative Color Doppler image of middle cerebral artery (MCA) arising from internal carotid artery (ICA), a typical doppler wave, and formulas used to calculate resistance and pulsatility indexes in MCA. Panel B depicts MCA velocity measurements. Panel C shows data for pulsatility index (PI) and resistance index (RI) of MCA. Data are mean ± SEM, n=10-19.

Cognitive function was assessed at 2M using touchscreen-fitted operant chambers. TGA-PE rats required more sessions and total trials to acquire the pairwise discrimination task (>80% accuracy for 2 consecutive days), and only 4 of 6 TGA-PE rats acquired the task within 40 days (Figure 5A). In the progressive ratio task, TGA-PE rats completed fewer ratios than SD rats, indicating reduced effort to obtain rewards (Figure 5B).

**Figure 5.**
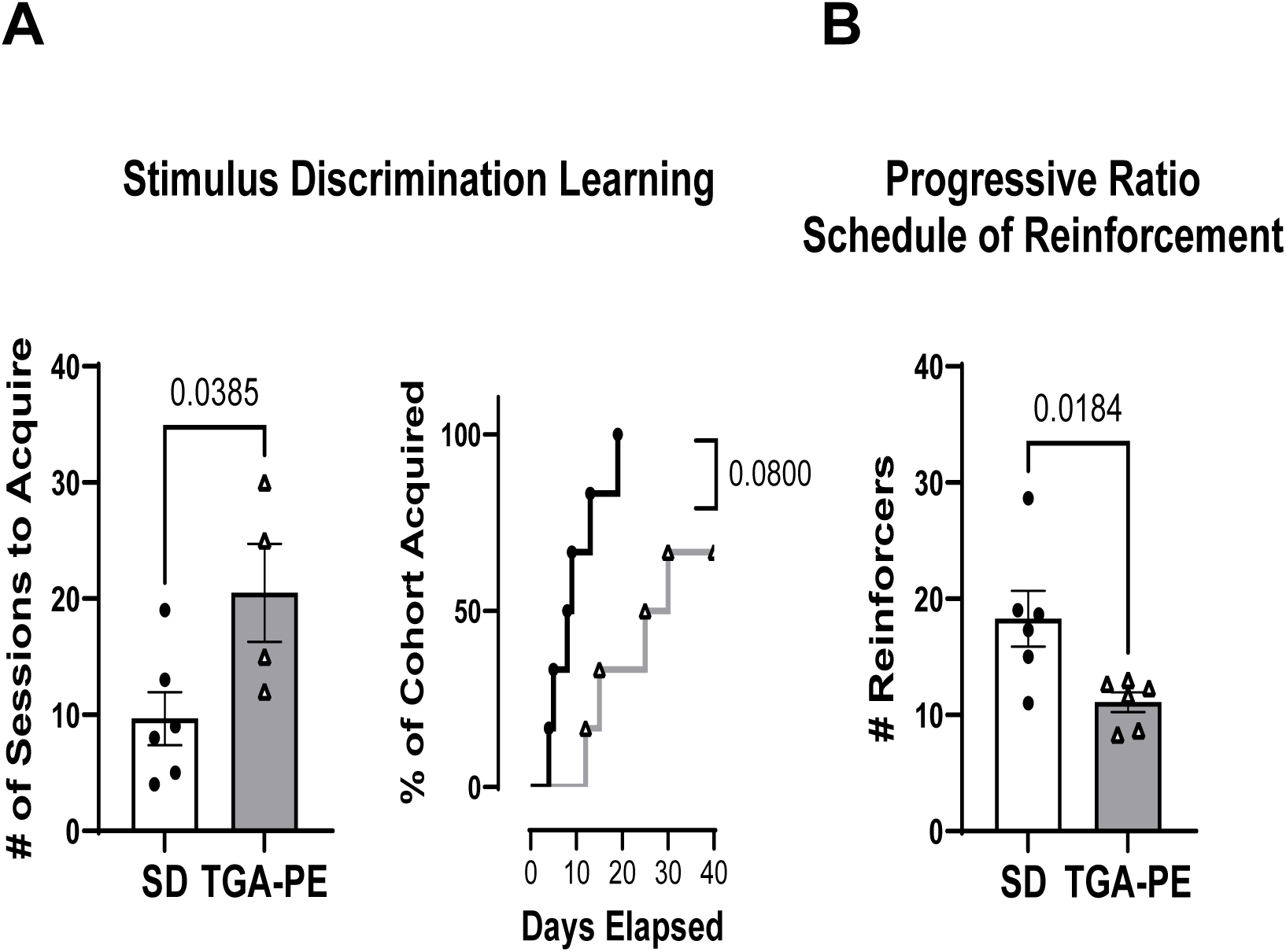
Cognitive assessments in TGA-PE vs. SD rats at 2M post-delivery. Panel A shows stimulus discrimination learning: number of sessions to reach the stimulus (left) and cumulative percentage of each cohort that reached the criterion (right) (Panel A). Only rats that reached the criteria were included. Panel B: Number of reinforcers earned under a progressive ratio schedule of reinforcement. Data are mean ± SEM, n=4-6.

Ultrasound-guided PAI showed no differences in MCA oxygenation or hemoglobin concentration in the temporal lobe between TGA-PE and SD rats at 2M (Figure 6A,C,F). In contrast, oxygenation of the temporal brain region adjacent to the MCA was higher in TGA-PE rats than in SD rats, without differences in hemoglobin concentration between groups (Figure 6B,E,F). To validate the PAI findings, MCA and brain sections were stained for the hypoxia marker HIF2α and VEGF. HIF2α levels did not differ in the MCA between TGA-PE and SD rats (Figure 6H) but were lower in the brain of TGA-PE rats than in SD rats (Figure 6J). No differences in VEGF levels were found in either MCA or adjacent brain areas (Figure 6I,K).

**Figure 6.**
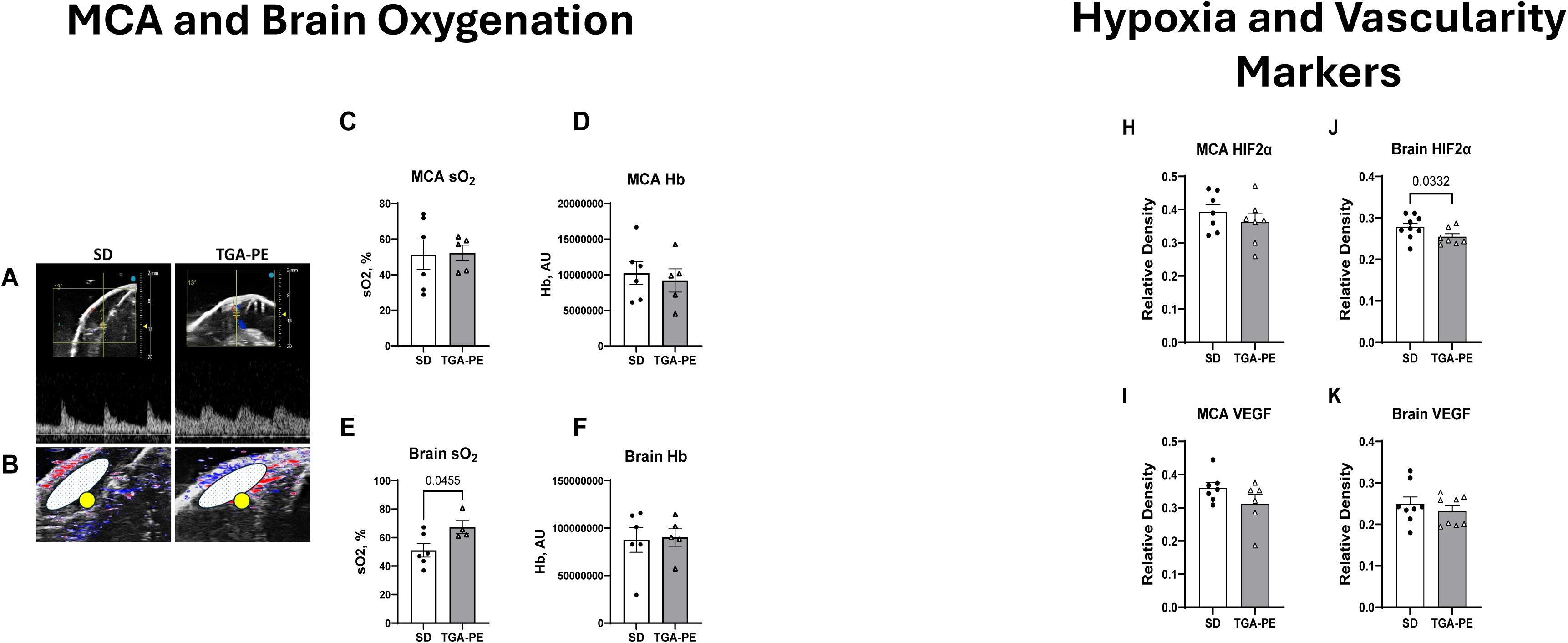
The PAI signal and quantification of oxygenation in MCA (Panels A,C,D) and adjacent temporal brain region (Panels B,E,F) determined by the photoacoustic imaging in SD vs. TGA-PE at 2 months post-delivery (2M). Panels H-K show the analysis of the immunohistochemistry for HIF2α (Panels H,I) and VEGF (Panels J,K) . Data are mean ± SEM, n=5-6. Abbreviations: AU, arbitrary units; sO2, oxygen saturation; Hb, hemoglobin.

To assess structural changes in the MCA, total collagen and elastin content were measured in TGA-PE and SD rats at 2M. Total collagen content did not differ between groups (Figure 7A), whereas total elastin content was lower in TGA-PE rats than in SD rats (Figure 7B). The collagen-to-elastin ratio did not differ between groups (Figure 7C).

**Figure 7.**
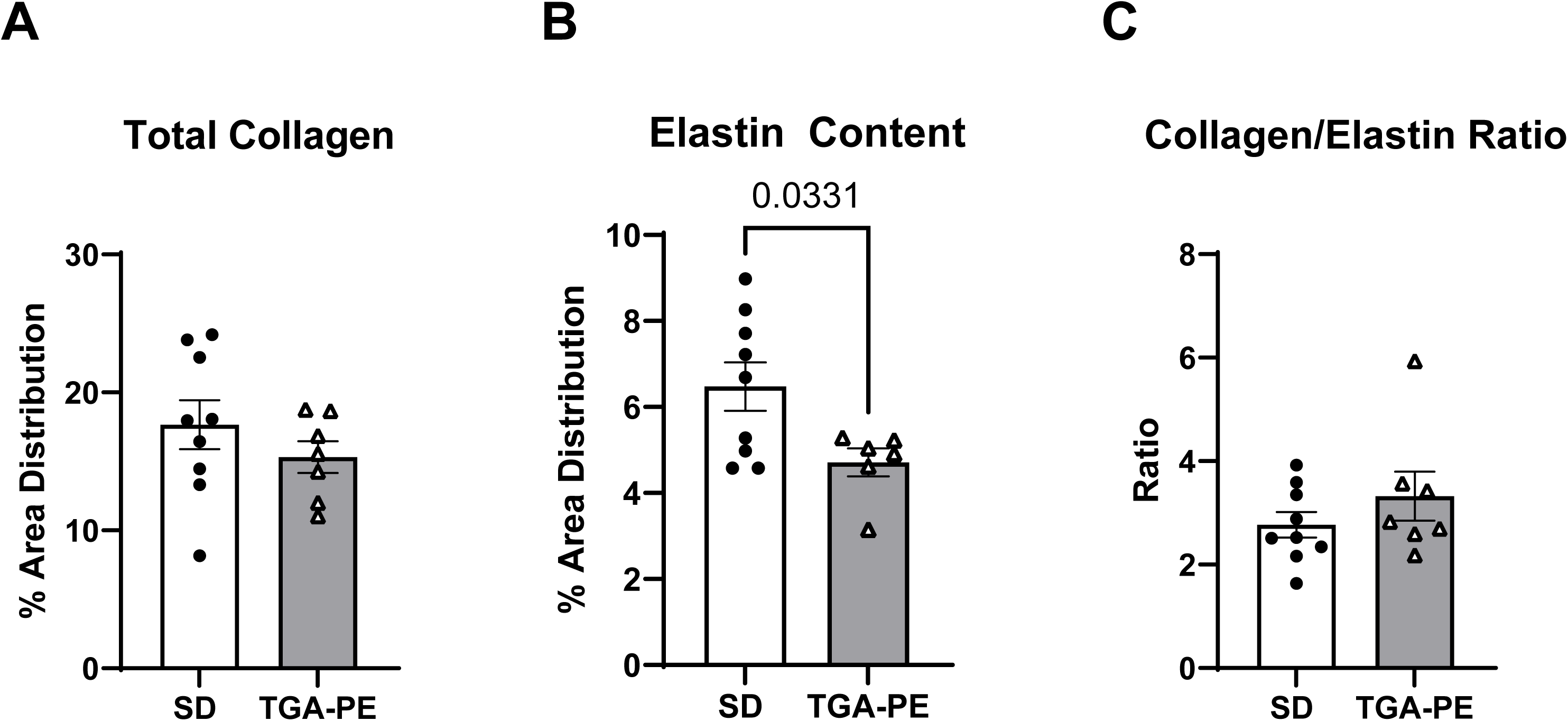
Total collagen and elastin content in MCA from TGA-PE and SD rats at 2 months post-delivery. Data are mean ± SEM, n=6-9.

Total EVs isolated from plasma were analyzed for size and concentration by nanoparticle tracking analysis, confirming a small EV size range of 120-160 nm. At late gestation, total EV size was smaller in TGA-PE rats than in SD rats. In SD rats, EV size was smaller at 2M than at late gestation, whereas EV size did not differ between late gestation and 2M in TGA-PE rats (Figure 8A). Total EV concentration did not differ between groups at either time point (Figure 8B). Nano-flow cytometry showed that 6.53% ± 0.24% of total EVs were PLAP-positive PD-EV, with no significant differences among groups (Figure 8C).

**Figure 8.**
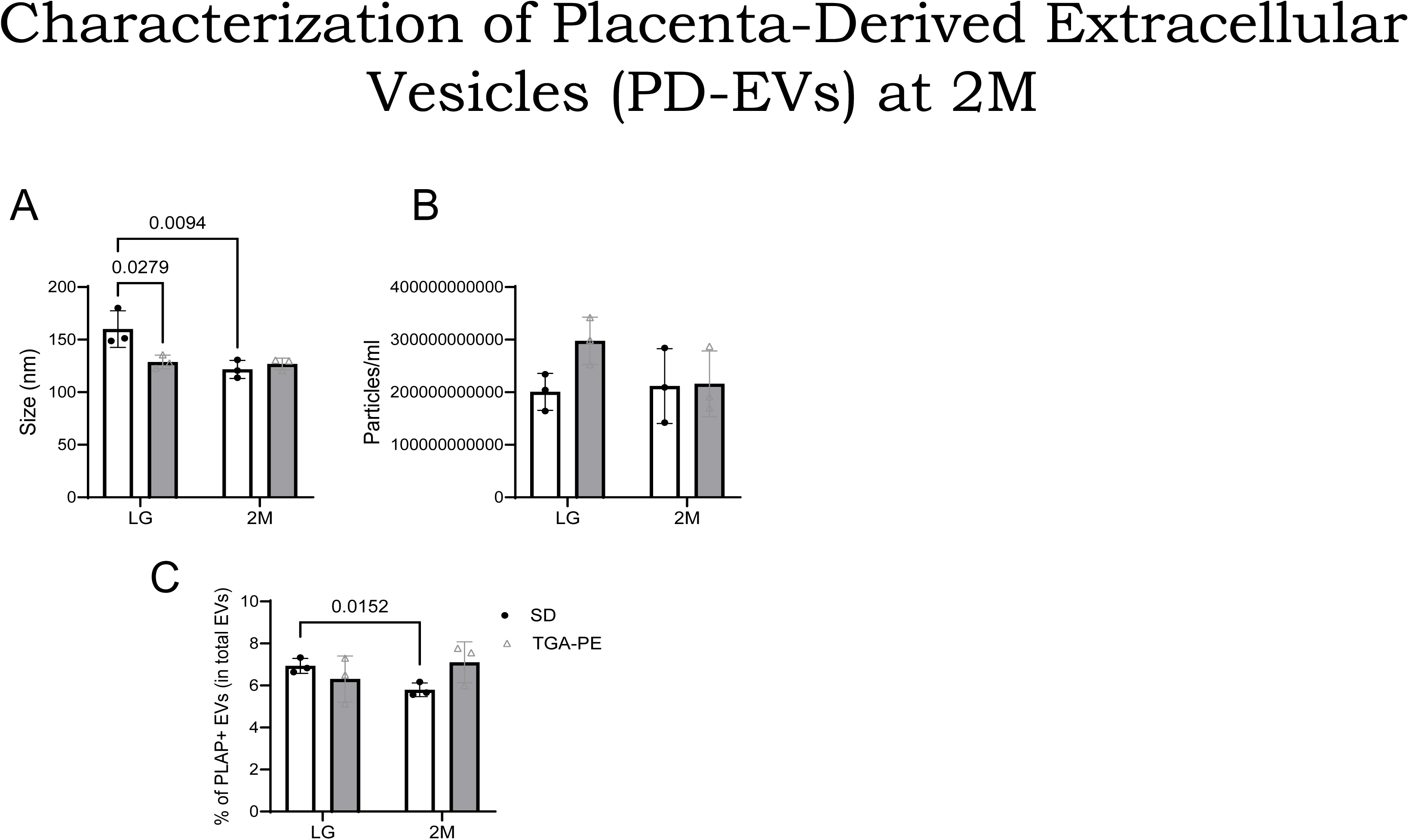
Total EVs, isolated from plasma, were first analyzed for size and concentration using nanoparticle tracking analysis in SD vs. TGA-PE at 2 months post-delivery (2M) (Panels A-B). Further total EVs were analyzed for the presence of PLAP+ EVs (PD-EV) using nano-flow cytometry and presented as percentage of PLAP+ EVs in total EVs (Panel C). Data are mean ± SEM, n=3 in each group.

PD-EV miRNA sequencing identified 29 of 2,436 miRNAs with significant differences between late gestation or 2M in TGA-PE rats compared with SD rats, including 17 upregulated and 12 downregulated miRNAs (Figure 9A,B). Validation of the sequencing results showed lower miR-363-3p levels in PD-EV from TGA-PE rats than in SD rats at 2M. Similarly, at 2M, miR-363-3p levels tend to decrease in PD-EVs (p<0.06; Panel C) and were lower in non-PDEVs and plasma from TGA-PE rats than in SD rats (Figure 9D, E). In HBEC-5i cerebral endothelial cells we found that the inhibition of miRNA363-3p with specific inhibitor upregulated the markers related to cellular senescence β-Gal and p21 (Figure 9F,G) and oxidative stress marker NOX-4 (Figure 9H).

**Figure 9.**
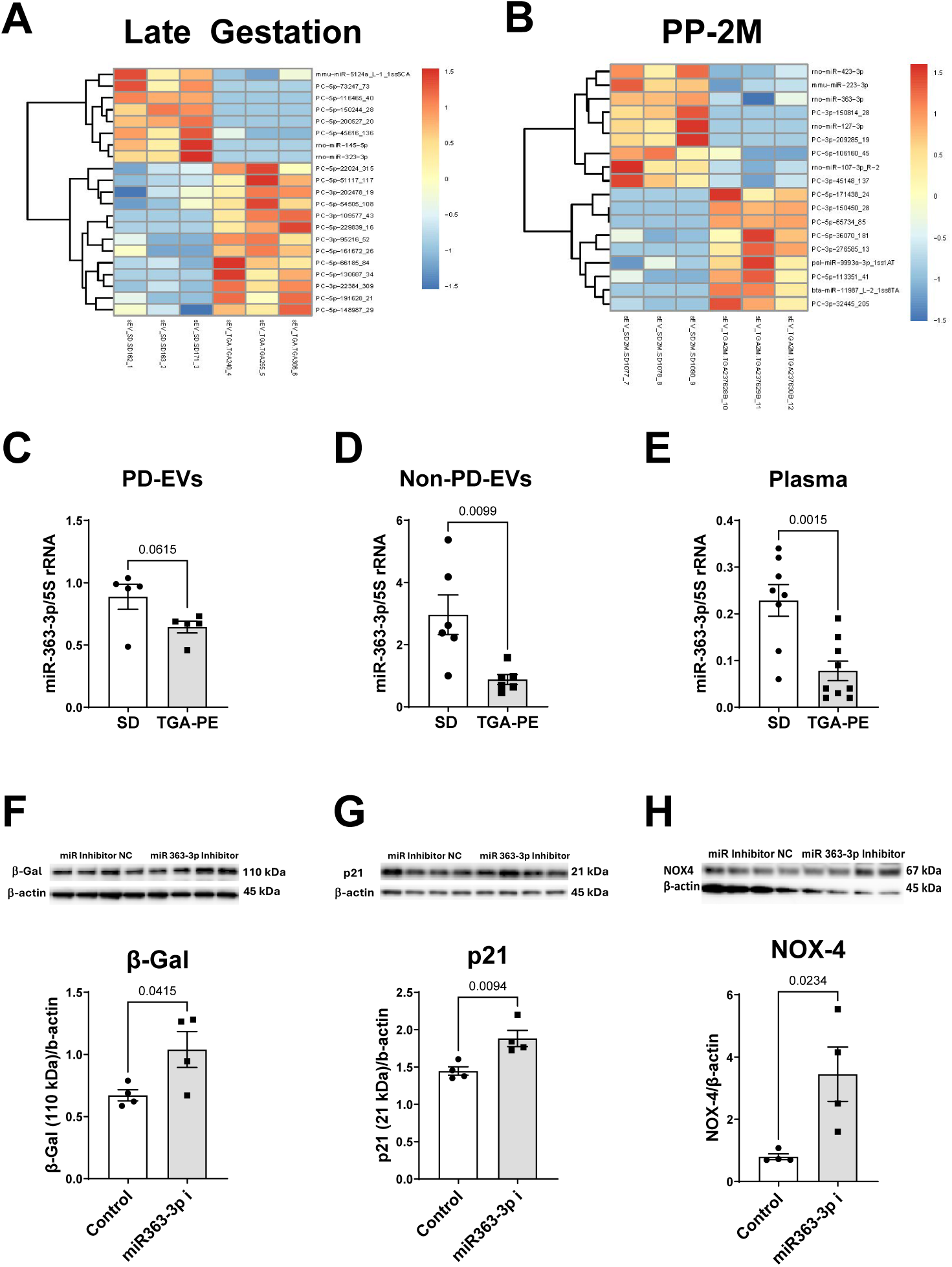
miRNA Seq of PD-EVs in SD vs. TGA-PE at Late Gestation (LG; Panel A) and 2 months post-delivery (2M; Panels B-E). The miRNA seq revealed significant alterations in LG (Panel A) or 2M (Panel B) for TGA-PE vs. SD. Middle panel demonstrates miRNA363 levels in PD-EVs (A), non-PD-EVs (B) or plasma (C). Panels F-G show changes in β-galactosidase (β-Gal), p21 (Panels F,G) and NOX-4 (panel H) in cultured cerebral endothelial cell line HBEC-5i. Data are mean ± SEM, n=6-9.

KEGG pathway enrichment analysis of 95BpPreg target genes identified seven pathways with elevated NEST scores: MAPK signaling, regulation of actin cytoskeleton, Ras signaling, lipid and atherosclerosis, cellular senescence, focal adhesion, and TNF signaling (Figure 10).

**Figure 10.**
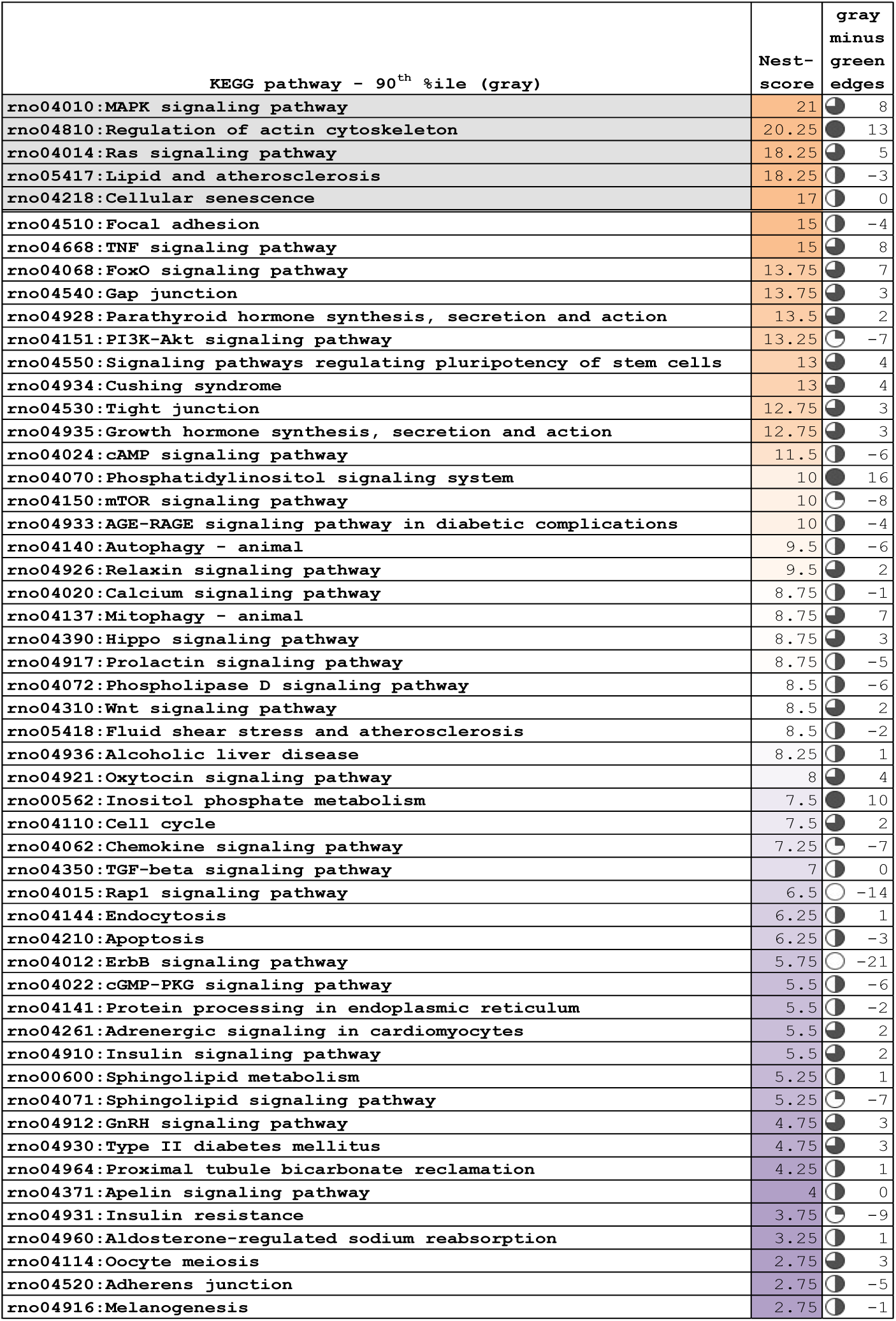
DAVID gene-enrichment analysis of KEGG pathway output by NEST-Score in miRNA from PD-EVs.

STRING-based network analysis [55] of the 39 MTS ≥ 95 target genes from these pathways, followed by Markov clustering, identified a primary cluster of 17 highly interconnected genes. Gene Ontology enrichment analysis of targets exceeding the upper inner fence with FDR < 0.05 identified pathways involved in vascular remodeling, cellular proliferation, migration, and angiogenesis, including epidermal growth factor receptor, vascular endothelial growth factor, transmembrane receptor protein tyrosine kinase, blood vessel morphogenesis, and angiogenesis. These GO terms were primarily associated with miRNAs that were downregulated in PD-EVs at late gestation and were upregulated at 2M in TGA-PE rats (Figure 11).

**Figure 11.**
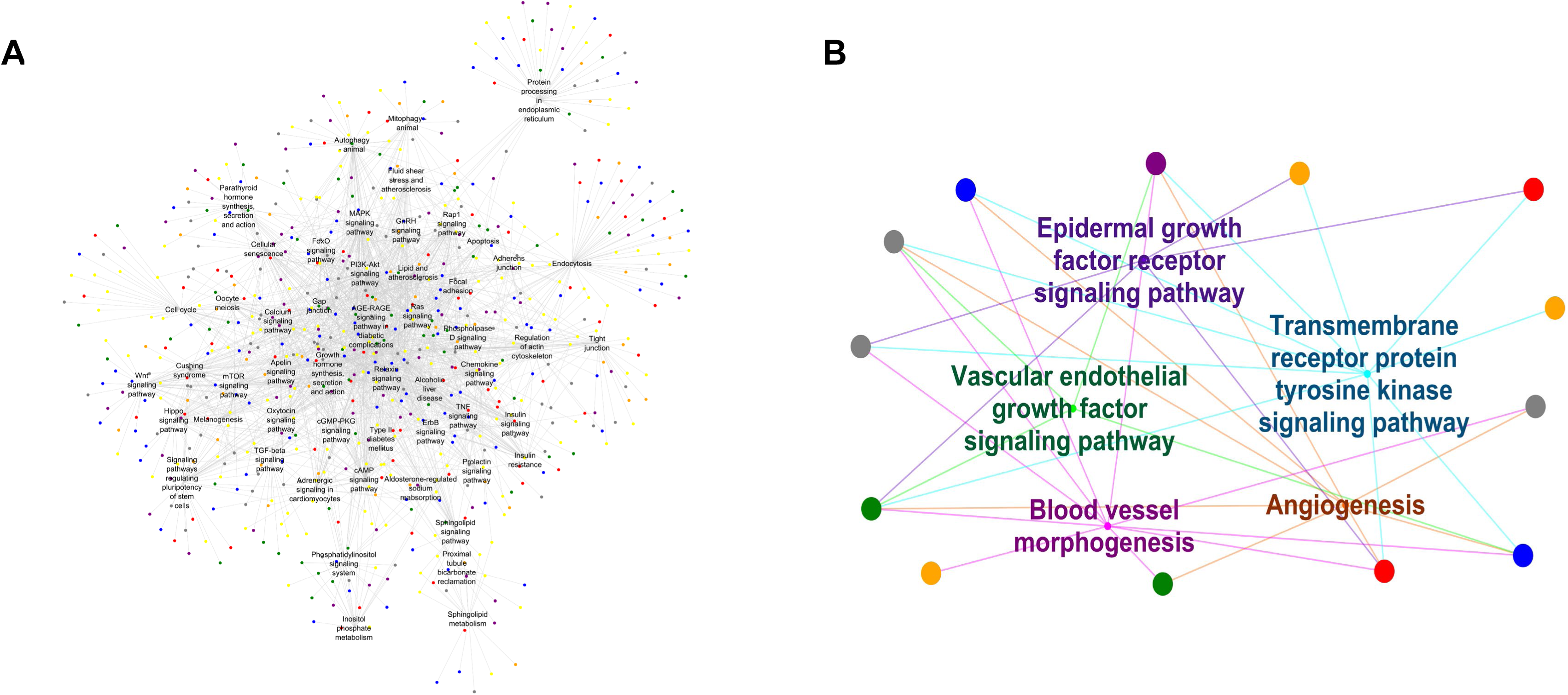
KEGG pathway/PD-EV miRNA target gene network (Panel A) and STRING-GO network (Panel B). Target gene color scheme is the same as described in Figure 1. Mapped in Cytoscape.

## DISCUSSION

Preeclampsia is increasingly recognized as a risk factor for long-term cerebrovascular disease and cognitive dysfunction. Clinical studies demonstrated that women with a history of preeclampsia have increased risk of chronic hypertension, stroke, cerebral small vessel disease, and cognitive dysfunction long after the index pregnancy, emphasizing the importance of identifying mechanisms that link pregnancy complications to lasting maternal cerebrovascular dysfunction [9]. Our findings demonstrate that at two months post-delivery TGA-PE rats exhibit hypertension, reduced MCA resistance and pulsatility indices with unchanged blood flow velocities, higher brain oxygen saturation with lower brain HIF-2α expression, reduced MCA elastin content, impaired associative learning and reduced reward-directed responding, and altered placental EV-derived miRNA profiles including systemically reduced miR-363-3p. Functional studies in human brain endothelial cells further showed that miR-363-3p inhibition upregulated markers related to oxidative stress and senescence. Together, these data support a model in which preeclampsia is associated with sustained maternal cerebrovascular and behavioral changes after pregnancy.

### Lower MCA resistance indexes

TGA-PE rats exhibited lower MCA resistance and pulsatility indexes at 2M despite higher blood pressure, whereas LMCA blood flow velocities were unchanged, suggesting changes in cerebral circulation. The two-month post-delivery timepoint in our study was selected to move beyond the immediate postpartum transition and assess persistent rather than immediate postpartum effects, a rationale supported by studies showing that vascular dysfunction persists years after preeclampsia. We confirmed significant differences in blood pressure between preeclamptic versus normal pregnancies at late gestation in TGA-PE [32]. Although the blood pressure decreased in preeclamptic model at 2M, it did not reach normotensive levels potentially suggesting an inverse relationship between MCA hemodynamics and systemic blood pressure. Higher heart rate also agreed with our previous reports suggesting autonomic dysfunction in this model [32]. Clinical evidence suggests that preeclamptic women exhibit impaired MCA function and cerebral hemodynamics. A lower MCA velocity was reported in postmenopausal women, while slightly higher MCA velocity was reported at 2-3 years post-pregnancy [7, 11, 56–58]. In addition, the response to CO_2_ was altered in preeclamptic women suggesting poorer autoregulation [59, 60]. Overall, the TGA-PE model provides a useful experimental system for investigating candidate mechanisms that may contribute to long-term maternal neurovascular changes following hypertensive pregnancy, although the extent to which these findings translate to human preeclampsia require continued investigation.

### Cognitive dysfunction

Because alterations in the cerebral vasculature have the potential to contribute to cognitive dysfunction, cognitive performance was evaluated at 2M. Our data showed that TGA-PE rats required more sessions and total trials to acquire the pairwise discrimination task, and fewer TGA-PE rats reached the acquisition criteria within 40 days. TGA-PE also completed fewer ratios in the progressive-ratio task, indicating reduced reward-directed responding. These behavioral findings were observed at the same time point when systolic and diastolic blood pressures remained elevated. The cognitive phenotype in TGA-PE rats is consistent with clinical studies reporting cognitive dysfunction months to years after preeclampsia, worse information-processing speed and executive function in midlife approximately 10.5 years after delivery and attenuated executive function or cognitive impairment decades after affected pregnancies [4, 6, 12, 13, 61]. Human data also link cognition to cerebrovascular changes. Reduced parietal cerebrovascular reactivity has been associated with poorer executive function and processing speed years after delivery in women with prior preeclampsia and placental maternal vascular malperfusion [12, 13].

Preclinical data support a vascular contribution to cognitive dysfunction, although maternal cognitive testing after experimental preeclampsia remains limited. During pregnancy, RUPP studies have shown impaired MCA myogenic responsiveness, loss of cerebral blood flow autoregulation, blood-brain barrier disruption, and cerebral edema [14–16]. At two months postpartum, placental ischemia was associated with cortical edema and neuroinflammation [16]. Our findings extend this literature by showing impaired learning acquisition and reduced reward-directed responding at 2M in TGA-PE rats in addition to reduced MCA resistance and pulsatility indexes and lower MCA elastin content. These data support future studies on testing whether persistent cerebrovascular dysregulation, placental signaling, or both contribute to post-preeclampsia cognitive outcomes. Furthermore, whether the activated renin-angiotensin system characteristic for this model has contributed to the neurovascular dysfunction remains to be established. Placental dysfunction and widespread maternal endothelial injury are major consequences of excessive pregnancy-specific RAS activation [62–65]. It is possible that RAS disbalance triggers placental and endothelial dysfunction that extends beyond delivery and contributes to later neurovascular vulnerability.

### Ultrasound-guided photoacoustic imaging

Human studies link preeclampsia with persistent cerebrovascular dysfunction, brain structural changes, altered cerebral oxygen metabolism, and cognitive impairment [5, 11–13, 22]. MRI-based studies have reported lower brain volumes after hypertensive pregnancy, reduced cerebrovascular reactivity during breath-hold challenge, and associations between reduced cerebrovascular reactivity and poorer processing speed or executive function in women with prior preeclampsia [5, 12, 13]. Maternal oxygenation-focused studies using MRI-based oxygen extraction fraction mapping or near-infrared cerebral oximetry also suggest altered cerebral oxygen metabolism or oxygen saturation in preeclampsia [6, 57, 58]. However, these approaches did not evaluate MCA-adjacent cerebral oxygenation with MCA hemodynamics in the same experimental model. Photoacoustic imaging combines optical contrast with ultrasound spatial resolution to estimate tissue oxygenation *in vivo*. PAI detects endogenous chromophores, including hemoglobin. Blood oxygen saturation (sO₂) is calculated as the fraction of oxygenated hemoglobin relative to total hemoglobin based on the distinct photoacoustic spectra of oxygenated and deoxygenated hemoglobin. Total hemoglobin concentration (HbT), determined by PAI, is often used as an indirect measure of regional blood volume and may reflect vascular characteristics within the imaged tissue. Although HbT does not directly quantify vascular density or vascularization, previous studies have reported positive correlations between PAI-derived HbT and CD31-positive vascular staining on the periphery [66].

Preclinical studies using the RUPP model have demonstrated impaired cerebral blood flow autoregulation using in situ laser Doppler measurements [15]. Broader rodent brain PAI studies demonstrated that photoacoustic approaches could measure cerebral hemoglobin content and oxygen saturation [67, 68]. Therefore, the present study extends the application of ultrasound-guided PAI to the evaluation of MCA and MCA-adjacent cerebral oxygenation in a post-preeclampsia model. In our study, MCA oxygenation and total hemoglobin concentration were not different between groups, whereas temporal lobe oxygen saturation was higher in TGA-PE rats despite unchanged HbT suggesting increased tissue perfusion. Lower brain HIF-2α levels were consistent with the higher regional oxygen saturation measured by PAI. However, additional injury markers and longer follow-up are needed to determine whether these oxygenation changes precede structural injury, brain remodeling, or cognitive decline. Although the underlying mechanisms cannot be determined from the present study, altered oxygen delivery, oxygen utilization, and/or neurovascular coupling may contribute to the observed phenotype. The absence of differences in HbT and immunostaining for VEGF suggests that the increase in brain oxygen saturation is not potentially associated with detectable changes in regional blood volume or angiogenic signaling. Overall, PAI complements TCD-based MCA hemodynamic assessment by providing regional oxygenation information. It is possible that lower resistance may lead to the exposure of cerebral microcirculation to hyperperfusion resulting in blood brain barrier damage and local ischemic neuronal degradation and cognitive dysfunction [8]. However, the pathways involved in cerebrovascular dysfunction are not well understood.

### miRNA profiling, miRNA363-3p and markers of vascular injury

MicroRNAs regulate gene expression and intercellular communication at the maternal-fetal interface and have emerged as biomarkers and mediators of pregnancy disorders, including preeclampsia and fetal growth restriction [69, 70]. Because placental dysfunction and placenta-derived EVs contribute to the pathophysiology of preeclampsia we profiled miRNAs and target pathways in PD-EVs during late gestation and post-pregnancy at 2M. Of 2,436 miRNAs profiled, 29 showed significant differences in TGA-PE compared with SD rats at late gestation or 2M including 17 upregulated and 12 downregulated, suggesting persistent changes in placental EV miRNA cargo after pregnancy. Notably, total EV size was smaller in TGA-PE rats than in SD rats during late gestation but did not differ at 2M suggesting that placental alterations may alter EV biogenesis or release during gestation without a long-term effect on vesicle size post-pregnancy.

Among identified miRNAs, miR-363-3p was lower in PD-EVs, and significantly in reduced non-placental EVs, and plasma. miR-363-3p has been reported to decline in maternal blood weeks before clinical preeclampsia, including at 28 weeks of gestation and approximately 10–12 weeks before diagnosis [71]. Prior studies also link miR-363-3p dysregulation to impaired trophoblast invasion and spiral artery remodeling [72], reduced endothelial apoptosis, oxidative stress, and inflammatory signaling through NOX4-dependent pathways [73], and improved neurological outcomes in a post-stroke depression model [74] suggesting a protective role of miRNA363-3p in vasculature and cognition. However, the expression and the role of miRNA363-3p during normal pregnancy, PE or long-term postpartum are not yet defined. Consistent with above mentioned reports, miR-363-3p was lower in TGAPE-derived PD-EVs compared with SD controls at 2M, a time point comparable to approximately 4–5 human years after delivery. While not establishing the causality, these data suggest that reduced placental EV-derived miR-363-3p may be associated with pathways related to endothelial adaptation, oxidative stress, inflammatory signaling, and persistent maternal vascular or behavioral changes after preeclampsia. Importantly, miR-363-3p levels were significantly reduced in the non-placental EV fraction and plasma of TGA-PE rats, suggesting that the reduction in miR-363-3p may reflect long-term systemic alterations in maternal vascular, immune, or other organ-derived signaling beyond the placenta. While the TGA-PE model reproduces several important features of human preeclampsia, additional studies are required to determine whether EV-associated miRNA pathways contribute to the increased long-term cerebrovascular and cognitive risk observed in women with prior preeclampsia.

In human brain endothelial cells, the inhibition of miR-363-3p upregulated NOX4, p21, and senescence-associated β-galactosidase, suggesting a possible protective role for miRNA363-3p. However, whether NOX4 is a direct target of miR-363-3p or mediates the senescence phenotype through an intermediate pathway remains to be established. Because endothelial senescence and oxidative stress contribute to cerebrovascular aging and vascular remodeling, reduced miR-363-3p levels may represent a potential mechanism contributing to long-term maternal cerebrovascular vulnerability after preeclampsia.

### miRNA targets and pathways

NEST-Scoring identified enrichment of miRNAs target pathways involved in MAPK, Ras, TNF, focal adhesion, cytoskeletal regulation, and cellular senescence. These pathways are relevant to endothelial dysfunction, inflammatory signaling, angiogenic regulation, and vascular remodeling. Among predicted miRNA targets, KLF2, NOX4, PTEN, and the PEDF/VEGF axis are biologically relevant to vascular health and remodeling. Since these targets were identified through bioinformatic prediction and pathway enrichment; direct validation in cerebrovascular tissue is needed to establish whether they mediate the postpartum phenotype.

Collectively, the miRNA target analysis supports a framework in which altered placental EV-derived miRNA cargo may be accompanied with changes in cerebrovascular function through pathways related to inflammation, angiogenesis, cytoskeletal organization, oxidative stress, and vascular remodeling. These findings also support the previously developed concept of the maternal brain at risk after preeclampsia [7]. The postpartum MCA hemodynamic and behavioral changes observed in TGA-PE rats may reflect neurovascular vulnerability associated with placental dysfunction during pregnancy. Importantly, network and gene ontology analyses further highlighted biological processes involved in angiogenesis, vascular morphogenesis, and cell proliferation. The observed temporal shifts in miRNA expression between late gestation and 2M suggest dynamic regulation of these pathways, potentially reflecting both acute and long-term vascular changes. Additional mechanistic studies will be necessary to determine whether specific EV-derived miRNAs directly drive these long-term maternal outcomes and whether they can be targeted therapeutically.

### Study limitations

Several limitations of this study should be acknowledged. First, the sample size was modest, which may have limited power to detect additional differences, particularly in the miRNA analyses. Second, our findings are correlative and do not establish causality between altered placental EV-derived miRNAs and longterm cerebrovascular or cognitive dysfunction. Third, only a limited number of candidate miRNAs were validated following sequencing; therefore, broader confirmation of the EV-PD miRNA profile will be necessary.

## Conclusion

Collectively, our findings demonstrate cerebrovascular, cognitive, and behavioral alterations in TGA-PE rats two months post-delivery, including hypertension, reduced MCA resistance and pulsatility indexes, decreased cerebrovascular elastin content, altered miRNA profiles, impaired learning, and altered reward-related behavior consistent with an anhedonia-like phenotype. These findings align with the emerging concept of a “maternal brain at risk” after preeclampsia and with clinical evidence of increased long-term cerebrovascular and cognitive vulnerability following preeclampsia [4, 7, 75]. The novel NEST-Scoring analysis provides an alternative approach for investigating alterations in biological pathway networks associated with these outcomes. Future studies are needed to determine whether the observed miRNA alterations contribute to vascular and cognitive dysfunction or serve as biomarkers of long-term maternal health outcomes. Overall, these findings provide new insight into cerebrovascular and neurobehavioral alterations following preeclampsia and highlight the need to further define the mechanisms linking pregnancy complications to long-term maternal brain health.

## Supporting information

suppl_plate

## Acknowledgements

We acknowledge the assistance of the Wake Forest University School of Medicine Preclinical Ultrasound and Photoacoustic Imaging Core and the Hypertension and Vascular Research Center Transgenic Core.

## Data availability statement

The data that support the findings of this study are available from the corresponding author upon reasonable request.

## Ethics Approval Statement

The Institutional Animal Care and Use Committee of Wake Forest University School of Medicine approved the study (A21-057), and all experiments were conducted in accordance with relevant guidelines and regulations.

## AI Technology Disclosure

The authors acknowledge the use of Copilot Claude 5.5 (accessed September 2026) to support creation of the graphical abstract figure. All AI-generated suggestions were reviewed, revised, and approved by the authors, who take full responsibility for the accuracy and integrity of the work.

## Author Contributions

Jonathan Ray - Study Design, Literature Search / Data Acquisition, Data Analysis and Interpretation, Manuscript Drafting;

Brian M. Westwood - Study Design, Literature Search / Data Acquisition, Data Analysis and Interpretation, Manuscript Drafting, Critical Revision;

Xuming Sun - Data Acquisition, Data Analysis and Interpretation, Manuscript Drafting, Critical Revision;

Robert Gould - Study Design, Literature Search / Data Acquisition, Data Analysis and Interpretation, Manuscript Drafting, Critical Revision;

Yixin Su - Data Acquisition, Data Analysis and Interpretation, Manuscript Drafting;

Marli Graves - Data Acquisition, Data Analysis and Interpretation;

Nildris Cruz-Diaz - Data Acquisition, Data Analysis and Interpretation, Manuscript Drafting;

Sangeeta Singh - Data Acquisition, Data Analysis and Interpretation, Manuscript Drafting;

Ashish Kumar - Data Acquisition, Data Analysis and Interpretation, Manuscript Drafting;

Gagan Deep - Study Design, Data Analysis and Interpretation, Manuscript Drafting, Critical Revision;

Hossam A. Shaltout - Data Interpretation, Manuscript Drafting, Critical Revision;

NyKea L. Lewis - Literature Search;

Tao Ma - Manuscript Drafting; Critical Revision;

Jeffrey Denney - Manuscript Drafting, Critical Revision;

Victor M. Pulgar - Manuscript Drafting, Critical Revision;

Liliya M. Yamaleyeva - Concept and Design, Literature Search / Data Acquisition, Data Analysis and Interpretation, Manuscript Drafting, Critical Revision, Supervision & Funding Acquisition

---

**Supplemental Figure 1.**
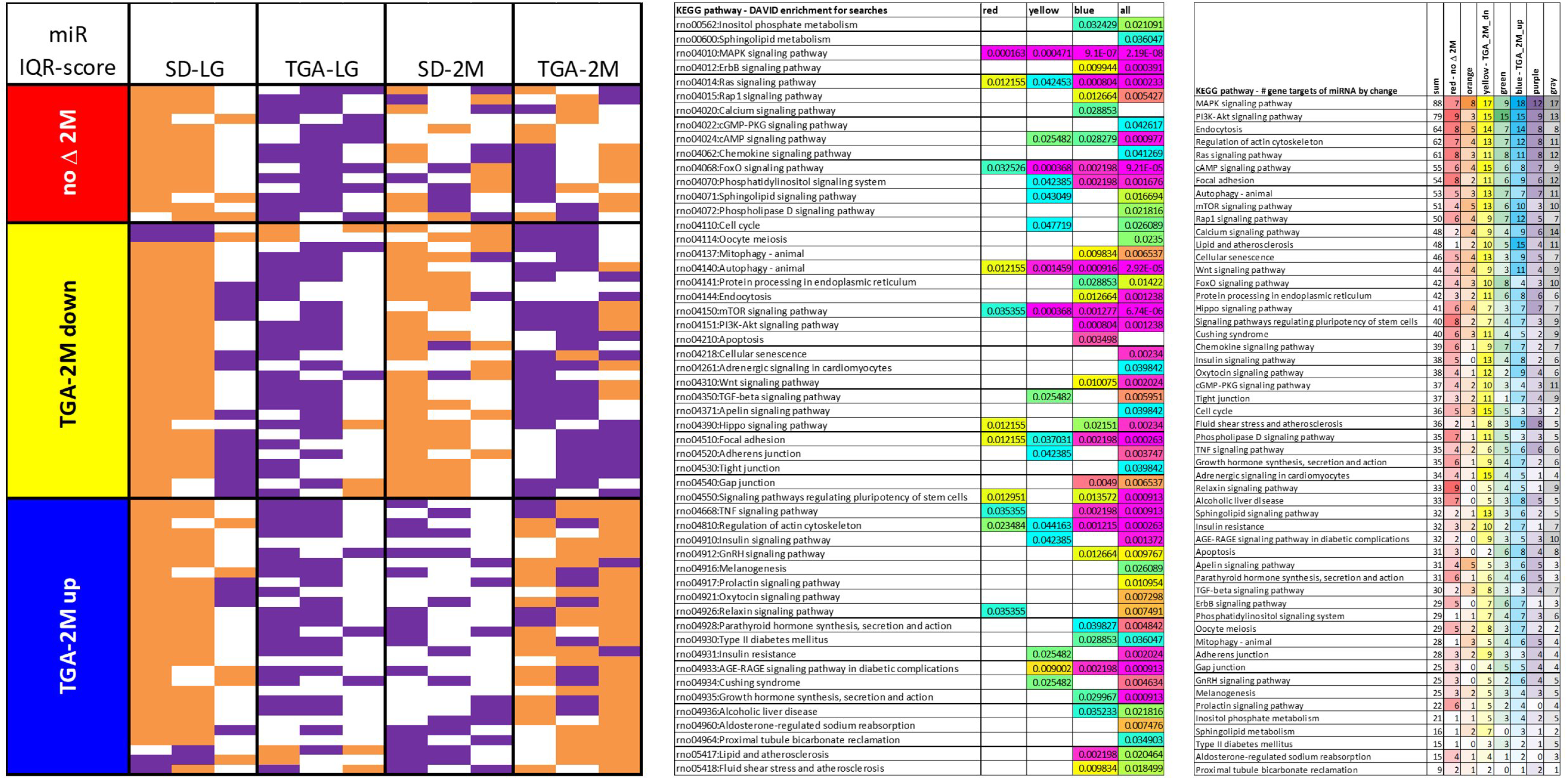
53 KEGG metabolic pathways scored as outlined in NEST-Scoring method and Figure 2.

**Supplemental Figure 2.**
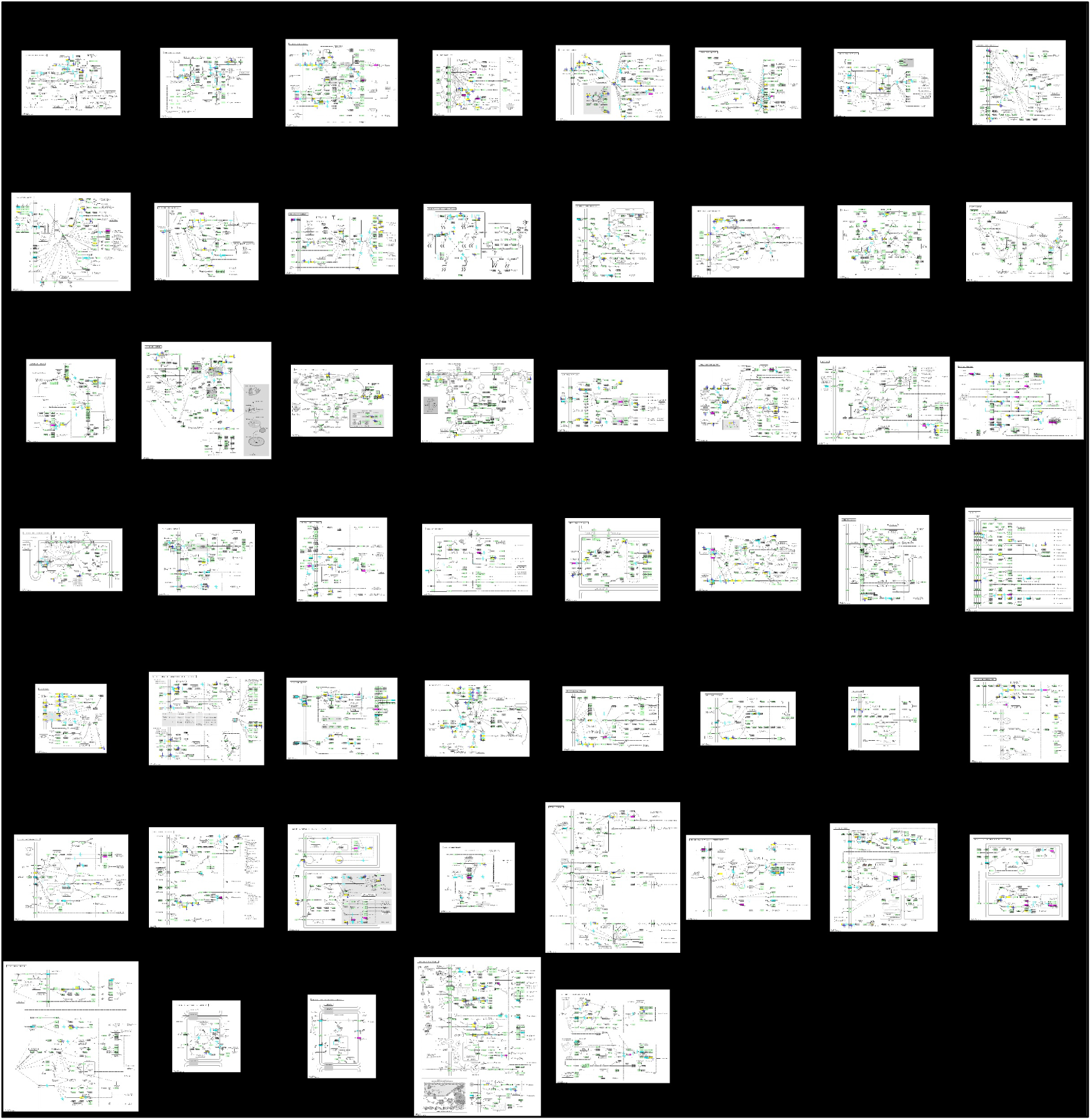
On the left, 70 miRNA separated into the three DAVID target search groups (individual subject [column] detail view from Figure 1, upper left) showing IQR-score inputs: +1 [orange], -1 [purple] and 0 [white]; each row is an individual miRNA. Middle table is the DAVID enrichment score outputs for each individual and the composite search. On the right, delineates where the target genes from each KEGG pathway fall into the Figure 1 triangular compositional diagram color schema.

