## Supplementary figures and images for "Long-Term Neurovascular Dysfunction and Altered Placenta-Derived EV miRNA Profiles in a Rat Model with Preeclamptic Features"

### suppl_table

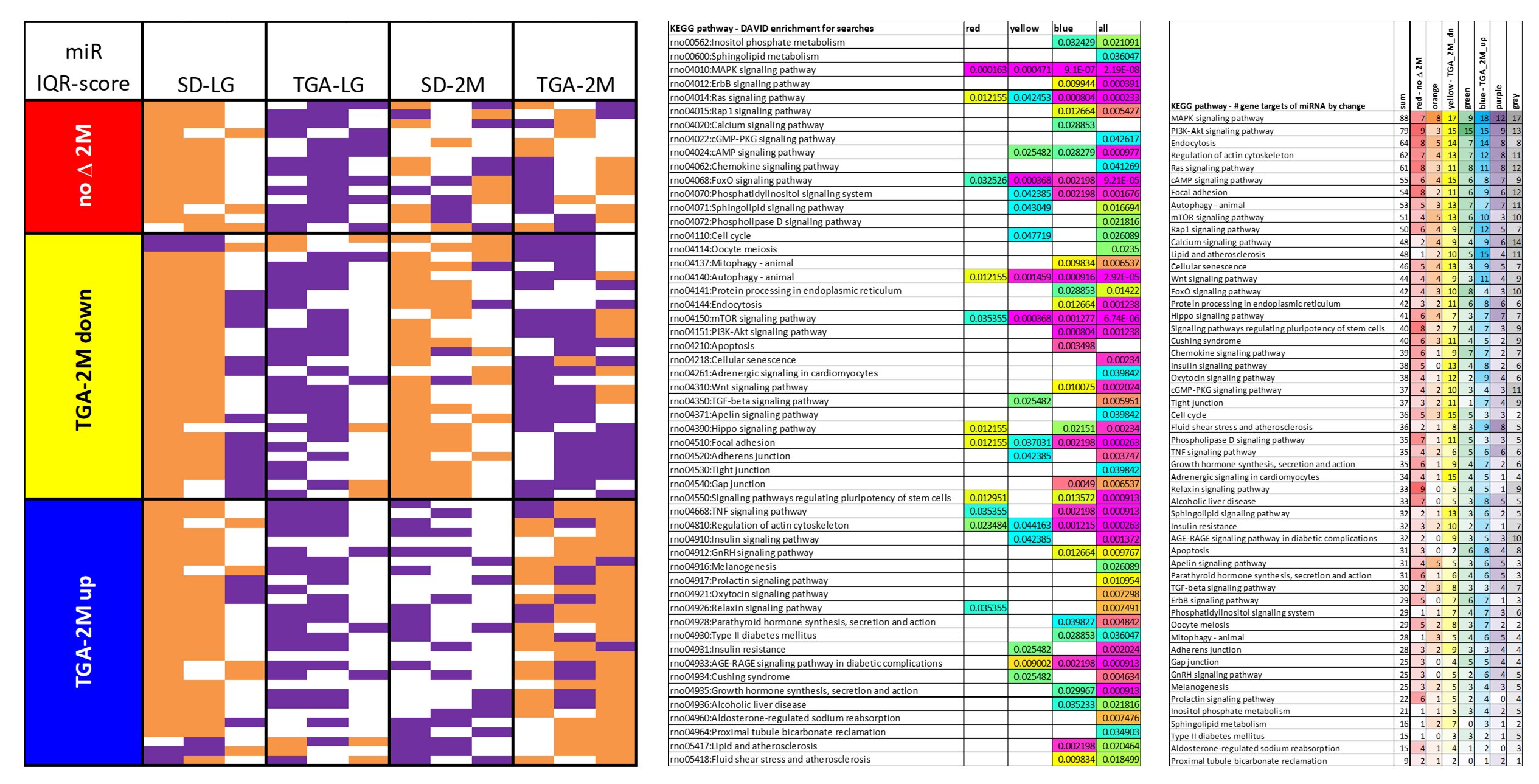
